# Beta burst waveform extraction using novel ^4^He-OPMs

**DOI:** 10.1101/2025.07.10.663870

**Authors:** Tjerk P. Gutteling, Maciej J. Szul, Sébastien Daligault, Etienne Labyt, Julien Jung, James J. Bonaiuto, Denis Schwartz

## Abstract

Studying the electrophysiology of motor preparation and execution is challenging due to the restrictions often placed on the experimental paradigm by the imaging modality. MEG is well suited to track temporal brain dynamics while offering good spatial resolution, but requires an absence of head motion due to the fixed helmet and associated cryogenic cooling system. Here we used novel, room temperature, wearable optically pumped magnetometers using Helium to measure the MEG in a motor preparation and execution task and compare this with ‘classic’ SQUID-MEG. Beta band activity (13-30Hz) is widely associated with motor activity, and it has become widely accepted that beta activity occurs in bursts rather than sustained activity. Here we used a novel beta burst extraction pipeline to look at not only the occurrence of the beta bursts across the timecourse of motor preparation and execution, but at the specific waveforms of the beta burst that evolve over time. Results show that specific beta burst waveforms show strong task relevant modulations of burst rate. Beta burst waveforms extracted using Helium-OPMs were very similar to those extracted using SQUID-MEG, with comparable modulations of burst rate over time for specific waveforms. This shows a promising step to obtaining high quality electrophysiological data in less restricted paradigms.

## 1. Introduction

Turning perception into action is one of the most important tasks our brains have to accomplish. While current neuroimaging methods are well-suited to studying perception, action is particularly challenging to study because most of these methods function optimally under conditions of minimal movement, and therefore severely hinder the range of experiments that can be performed. Moreover, the neural field signatures of action preparation and execution appear to be more nuanced than classically assumed, requiring precise, high-SNR recording even in the presence of movement.

Activity in the beta band (13-30 Hz), for example, has long been known to reduce before action onset and rebound above the baseline after action execution (Neuper et al., 2006; Pfurtscheller and Lopes da Silva, 1999). However, mounting evidence suggests that trial-averaged beta power is the result of probabilistically occurring transient bursts at the single trial level (Echeverria-Altuna et al., 2022; Jones, 2016; Sherman et al., 2016), which better explain variation in response time and accuracy than averaged beta power (Enz et al., 2021; Little et al., 2019). Additionally, there is evidence that the waveform shape is also behaviorally relevant (Higgins et al., 2022; Marshall et al., 2022; Quinn et al., 2021). While the average beta burst waveform has been consistently found to have a typical wavelet-like shape, a large amount of variability exists in the individual burst waveform shapes, and burst with different shapes have different rate dynamics (Papadopoulos et al., 2024; Rayson et al., 2023; Szul et al., 2023). The ideal neuroimaging method for studying action should therefore have high temporal and spatial resolution, robustly deal with subject movement, and be capable of acquiring high quality signals for analysis of burst waveforms.

Magneto-encephalography (MEG) is very well suited to record brain dynamics with high temporal- and good spatial resolution. However, classic SQUID (superconducting quantum interference device, Cohen, 1972) based MEG requires helium cooling and a fixed sensor array in a large rigid dewar, requiring an almost complete absence of head movement from participants. Alternatively, electro-encephalography (EEG; Berger, 1929; Biasiucci et al., 2019) involves wearable sensors, but suffers more from muscle artefacts, as well as poorer spatial accuracy and signal quality relative to MEG (Gross, 2019). Optically pumped magnetometers (OPMs) have recently made great advances and offer a solution to the restrictions of SQUID-MEG while retaining the benefits over EEG for motor task paradigms (Brookes et al., 2022). OPMs are wearable MEG sensors that register modulations in the magnetic field using a gas cell as the sensitive element and laser absorption to probe density fluctuations. Recent studies using OPMs have shown very promising results, obtaining good quality data under conditions of movement (Boto et al., 2018), recording beta-band bursts (Rivero et al., 2024), including in different age groups (Boto et al., 2022; Rier et al., 2024), and even whole-body movement (Holmes et al., 2023; Mellor et al., 2023; O’Neill et al., 2025; Seymour et al., 2021). These studies highlight the success of OPM applications in situations unsuitable or suboptimal for cryogenic MEG. The majority of OPM studies have used sensors based on alkali gas (usually Rubidium), which have excellent sensitivity in the range of 7-15 ft / √Hz, but suffer from a limited dynamic range (∼± 5 nT) (Allred et al., 2002; Tierney et al., 2019). This is especially problematic in paradigms with (head) movement and necessitates the use of advanced active shielding to obtain useable data.

The current study uses a prototype system of tri-axial OPMs based on Helium-4 as the sensitive gas element (^4^He-OPMs, MAG^4^Health, Grenoble, France) (Beato et al., 2018; Labyt et al., 2019). The key benefit of their construction is the large dynamic range (>200 nT) due to a closed loop operation in combination with a large resonance bandwidth. This allows for much larger head (sensor) movements without saturation of the signal, in environments with modest shielding. This comes at the cost of sensitivity, which is around 45 ft / √Hz or better on two of the three measurement axes for the sensors used in this study (Fourcault et al., 2021; Gutteling et al., 2023). However, these sensors operate at room temperature due to the use of Helium gas and thus do not need any insulation (alkali OPMs operate at ∼150 °C), allowing placement directly on the scalp, thus increasing SNR and providing some compensation for the reduced sensitivity (Iivanainen et al., 2017). ^4^He-OPMs therefore potentially offer the ideal combination of dynamic sensor range and increased SNR for studying waveform-specific burst dynamics during motor tasks.

To gain a first evaluation of this prototype system in a motor task, we use a reaching task with moderate movement in a standard shielded room. We use a novel burst extraction pipeline, previously described in Szul *et al*. (2023) to extract beta bursts from classic SQUID-MEG and ^4^He-OPM data during a simple motor task to assess the potential of wearable MEG for the application of motor tasks.

## 2. Methods

### 2.1 Participants

A total of 18 healthy participants (age range 19-51, mean age 33.1 ± 8.6 years, 2 were left-handed) took part in the current experiment as part of a larger study with multiple tasks (Gutteling et al., 2023). They had no history of neurological or psychiatric disorders and were not taking any medication active in the central nervous system. All participants signed an informed consent form prior to participation. The study was approved by the regulatory and ethics administration in France (IDRCB nr. 2020-A01830-39).

### 2.2 Task

Participants performed a cued visuomotor reaching task using a joystick (NATA technologies, Canada), which was a simplified version of the task described in Szul *et al*. (2023), see Figure 1. Each trial started with a central fixation target (0.6 × 0.6 degrees), shaped as a bullseye with crosshair (Thaler et al., 2013). After a variable delay (1-2s) a circular random dot kinematogram (RDK; 7° radius, 200 white dots, circular movement) appeared for 2 seconds. The motion was partially coherent, but this was not relevant for the current analyses. Along with the RDK, five circular targets appeared on the top outer edge of the dot pattern, spaced 30° apart between the central midline (0°) and 120° eccentricity (clockwise). One of these targets was larger (3.25°) and green, which indicated this target as the reach target for this trial. The other targets were smaller (1.625°) and gray. After the offset of the dot motion, all targets remained on screen for a variable delay period (0.5 -1 s) after which the gray targets disappeared and the green reach target remained. This served as the ‘go’ cue for the participant to move the joystick, controlling a small 0.5° white square, towards the reach target. The trial ended when the white square reached the edge of where the RDK was (7° from center). Feedback was then given on the final position of the cursor as a red dot, which was on screen for 2s. Trials with movements before the go-cue or late movements (>1s) were discarded. After the feedback there was a variable 1.5-2 s interval where the participants were to bring the white square back to the central fixation point.

**Figure 1.**
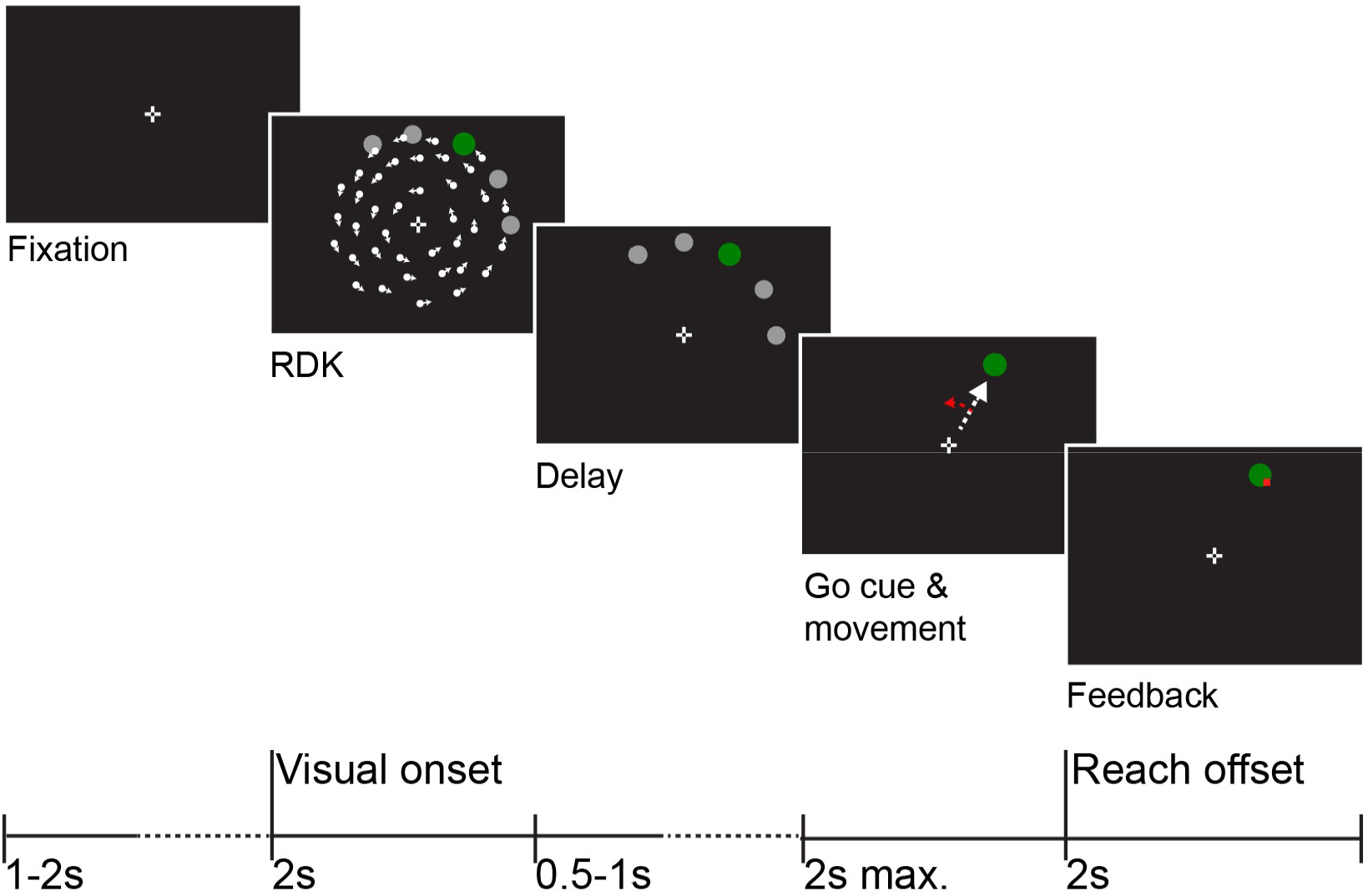
Task overview. Participants performed a cued reaching task. After fixation, a random dot kinematogram (RDK) was shown along with several targets of which one was the reaching target. This target was green and larger. This is the ‘visual onset’. After a variable delay, the other targets disappeared, which was the go-cue to start the reaching movement (white arrow, not visible). During reaching, a motor rotation could be applied (red arrow), although the cursor was invisible. When the target was reached, or a location equidistant to the target, feedback was shown. This was the ‘reach offset’.

Participants underwent 1–3 brief training blocks consisting of 12 trials each, with no visuomotor perturbation. During these blocks, the position of the cursor was visible throughout the entire trial (online feedback). When the reach distance exceeded the center of the instructed target, the cursor turned red. A brief message was then displayed for 2 seconds, indicating if participants initiated the reach too early, failed to complete it quickly enough, or deviated too far from the target’s center. After the training blocks, subjects completed a single block of 56 trials with no visuomotor rotation. However, during this block and the following blocks, the cursor disappeared once its distance from the fixation target exceeded 1° and reappeared, turning red upon reaching the target (endpoint feedback). Subsequently, participants completed a block of 56 trials with visuomotor rotational perturbation. In these trials, the visual position of the cursor was perturbed by rotating it by -30°. Finally, subjects completed one washout block consisting of 56 trials without visuomotor perturbation.

The experiment was conducted inside a dimly lit magnetically shielded room (MSR) and stimuli were projected on a screen in front of the participant using a Propixx projector (VPIXX Technologies Inc., Canada). The task was implemented using Psychtoolbox (v3.0.16; Brainard (1997) running on Matlab R2017b (The Mathworks Inc., Natick, Massachusetts).

### 2.3 Data acquisition

SQUID-MEG and OPM-MEG data were collected in a standard magnetically shielded room (2 *μ*-metal, 1 copper layer, Vacuumschmelze, Hanau, Germany) without any active shielding. *SQUID-MEG* data were acquired using a 275-channel CTF MEG system (CTF MEG Neuro Innovations, Inc., Coquitlam, Canada). The data were digitized at 1.2 kHz with a 300Hz low-pass filter. *OPM-MEG* recordings were made using a set of five triaxial ^4^He-OPM sensors (MAG^4^Health, Grenoble, France) (Beato et al., 2018; Labyt et al., 2019), with one of the sensors serving as a reference sensor located on the top of the head, 10 cm above the scalp. The remaining four sensors were placed in an area approximately covering the left sensorimotor hand area (see Fig. 2). As no individualized placement was done, sensor slots were chosen to cover a wider area to allow for individual variation. For the two left-handed participants, sensors were placed over the right hemisphere, and the task was performed with the left hand. As the sensors operate at room temperature, the OPMs were placed directly on the scalp of the subjects in open-ended slots of a semi-rigid helmet with 96 possible sensor positions. The subjects were seated on the same chair, in the same upright position as the SQUID-MEG acquisition. A wooden frame surrounded the head of the participant and supported the cables, relieving the head of weight, as well as reducing cable movement. The head was not further restricted from movement, although subjects were asked to minimize any movement. No subject reported discomfort related to the weight of the helmet and sensors, although one third reported a restriction of movement due to the fixation to the wooden frame. The data were digitized at 11kHz.

**Figure 2.**
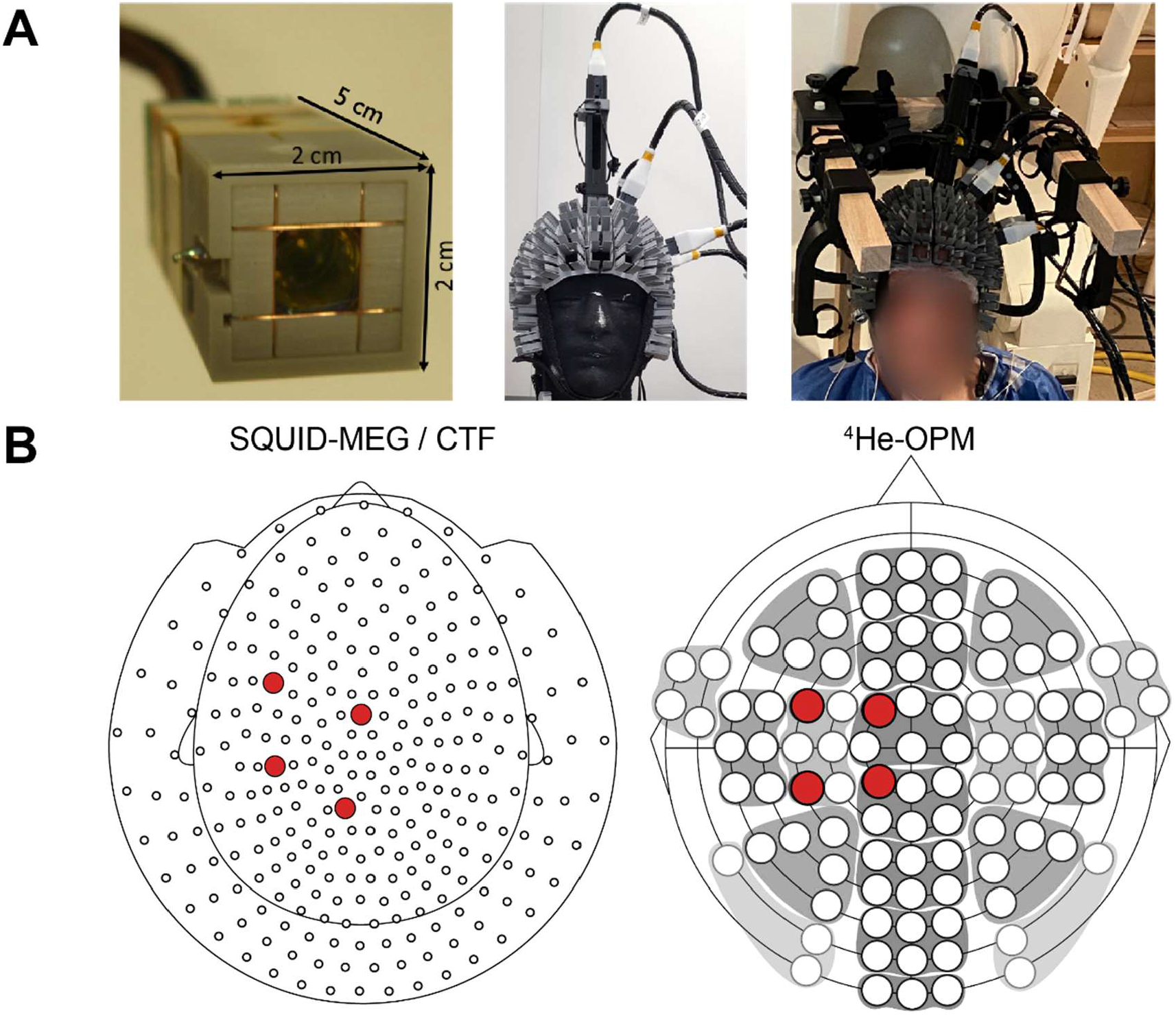
Experimental setup. A) left: The ^4^He-OPM sensor, viewed from the bottom, with the sensitive gas element to be placed on the scalp. Middle: four sensors were placed on the scalp and one was suspended 10cm above the head, mounted on a semi-flexible headcap. Right: the setup as used during the experiment. Cables were supported by a wooden frame. B) Sensors used in the analysis for SQUID-MEG (left) and OPM-MEG (right).

### 2.4 Data analysis

All electrophysiological data were analyzed using MNE-python (version 1.7.0, Gramfort et al., 2013) in a Linux environment. Analysis pipelines for SQUID-MEG and OPM-MEG data were kept as similar as possible to enable appropriate comparison, while optimizing data quality per modality. While the ^4^He-OPM are triaxial (one radial and two tangential axes), the sensitivity on the second tangential axis is approximately a factor 4 lower than the other axes (∼200 ft/√Hz vs. <45 ft/√Hz). We therefore chose to discard data from this axis. Before applying ICA decomposition (fastica), bad segments (spiking artifacts, high amplitude events) and breaks (>9 s) in the data were marked to be ignored, line noise (50Hz) and a video camera artefact (60Hz) was removed and the data were bandpass filtered between 1-100Hz. For the OPM data, due to the low sensor count, ICA components showing residual large amplitude noise were removed, which were usually confined to a single component. For SQUID-MEG ICA components related to eye movements, blinks, heartbeat and gross non-neural artifacts were selected to be removed. For SQUID-MEG 5.7 (± 2.5) components were removed and 1.05 (± .52) components for OPM-MEG. For an overview of the main results without the ICA step, see supplementary figures 8-10. After removal of these ICA components, further processing of the SQUID-MEG data was restricted to four sensors that were closest to the OPM sensor locations (see Fig. 2B). Sensor matching was done based on a prior digitization of the OPM helmet sensor positions using a Polhemus system (Polhemus, Colchester, USA). This was used as a reference for the OPM sensor positions to find the closest SQUID-MEG (CTF) sensor based on Euclidean distance. The data were then epoched either relative to the onset of the RDK visual stimulus [-1 s to 2 s, the ‘visual’ epochs] or relative to the end of the reaching movement [-1 s to 1.5 s, the ‘motor’ epochs]. For the SQUID-MEG dataset the electro-oculogram (EOG) was acquired, and used to regress out any (atypical) residual eye movement artefacts. The data were then downsampled to 1kHz. To further denoise the OPM data, the reference channel data from the sensor mounted 10cm on top of the head was used to remove residual low frequency noise. This was done by low-pass filtering the reference channel data at 10Hz to avoid introducing measurement noise from the reference sensors, and using linear regression per trial and sensor to remove the low frequency noise. Pilot studies have shown noise from the background field and subject movement is usually below 10-15Hz. As the reference sensor does not experience exactly the same background field and head movement as the other sensors, deflections can be shifted slightly in time. To accommodate this, dynamic time warping (dtw python package; Giorgino, 2009) was applied to the reference regressor to align with the individual sensor timecourses. This was applied if this resulted in a better fit, i.e. a lower root mean square (RMS) error between the unregressed signal and reference timecourse. The same procedure, including dynamic time warping, was used to remove the evoked component of the OPM data. An automated trial rejection procedure was used (autoreject package; Jas et al., 2017) to remove any artifactual epochs left in the data [SQUID-MEG: mean rejection ‘visual epochs’ 4.1% ± 7.3%; ‘motor epochs’ 6.7% ± 8.6%, OPM-MEG: mean rejection ‘visual epochs’ 17.1% ± 12.8%; ‘motor epochs’ 18.1% ± 14.5%].

### 2.5 Burst detection

For the detection of beta bursts, we used the iterative burst detection algorithm described in detail in Szul *et al*. (2023). In short, single trial time-frequency (TF) estimation was performed using the superlet transform (Moca et al., 2021), which aims to optimize the time and frequency resolution of the resulting TF estimation. We used an adaptive superlet transform based on Morlet wavelets with 2 cycles per frequency [2-100 Hz] and an order (the multiplication of the minimum cycle number) between 1 and 40. To check for significant modulation of beta band power, averaged and baselined [500-0 ms before visual onset] TF data were subjected to one-sample spatiotemporal clustering permutation tests over subjects (n=1000 permutations, cluster p<.05, 2-tailed). From the unaveraged data, the power spectral density per channel (sensor/axis) was extracted in the 15-45Hz range, which was used to estimate the aperiodic spectrum using the ‘specparam’ algorithm (Donoghue et al., 2020). This aperiodic spectrum was then subtracted from the single trial TF estimation. After this the largest peak in TF space (i.e. a burst) was iteratively fitted as a 2D Gaussian, yielding burst parameters (duration, amplitude, peak time, peak frequency, frequency span) and subtracted from the TF estimation. Only bursts in the 15-30Hz range were considered as ‘beta’-bursts. This process continued until there were no more maxima above the noise floor. The extracted burst peak times were then used to extract the burst waveforms from the epochs in the time domain (260 ms centered on the peak). To eliminate the influence of slower event-related dynamics on burst waveforms, we first regressed out the epoch-averaged signal from the timecourse of each trial. The sign of the extracted waveforms was flipped if the central deflection was not negative.

### 2.5 Burst analysis

As in Szul *et al*. (2023), all extracted burst waveforms were subjected to principal component analysis (PCA) to extract a range of waveform shapes for the beta bursts. To do this, first the extracted beta bursts were pooled together from all subjects, epochs, epoching conditions (visual/motor) and modalities (SQUID, OPM). The outside 10^th^ and 90^th^ percentiles of median amplitudes were removed as either high or low amplitude events are likely artefactual. The remaining burst waveforms (N = 699,507) were subjected to PCA analysis (20 components; PCA implemented in the scikit-learn toolbox, (v1.9.3, Pedregosa et al., 2012). Subjecting both modalities to a single PCA analysis ensures comparability of the resulting components and beta burst waveforms across modalities, as the data variability is divided along common axes. Each burst received a score for each of the 20 PCA components, representing the shape of the waveform along that dimension. For each component, the mean score was calculated over time by binning bursts in time (50ms bin width), and averaging their scores for that component, showing the modulation of this waveform component over time. Meaningful components, i.e. those that show modulation over time consistent with task performance, were selected based on changes in the mean score after the appearance of the visual cue, indicating waveform-specific rate modulation during the task performance. Specifically, the slope of the score over time for the visual epochs was calculated per component using a least-squares fit, to index modulation due to motor preparation, analogous to motor related beta suppression. We selected the three most modulated components between the two modalities (components 6,7 & 8, see Fig. 4B). The burst rate over time was calculated by binning bursts by time of occurrence (50ms bin width) and component score (40 bins based on score percentile), resulting in a burst rate over time for every component along the PC dimension. Baselined average burst rate, component specific burst rates and beta power timecourses were subjected to one sample (spatio-)temporal cluster permutation tests for test for significant deviations from baseline. Extracted burst parameters (burst duration, amplitude, peak frequency and frequency span) were compared between visual and motor epochs, as well as between modalities (SQUID-MEG, OPM-MEG) using Bonferroni-corrected Kolmogorov-Smirnov tests (Pratt and Gibbons, 1981).

### 2.5 SNR

To compare performance between SQUID and OPM-MEG, signal-to-noise ratios (SNR) were calculated for modulations in beta band power, as absolute maximum deflection relative to baseline divided by baseline standard deviation [500 to 250 ms before visual onset]. SNR values were tested using a repeated measures ANOVA with a single factor ‘sensor modality’ (SQUID, OPM_radial_, OPM_tangential_). Additionally, percent change from baseline was extracted per binned percentile for PCA components 6,7 and 8 and averaged 500-1500ms after visual stimulus onset for all sensor types. Average burst rate values were subtracted and the percentile bin with the largest percent baseline increase was chosen (the upper 75th-100th percentile in all cases). Resulting values were subjected to a repeated measures ANOVA with factors ‘PC component’ and ‘sensor modality’. Statistical analyses were made using the JASP software (v0.18.3, JASP team, 2024).

## 3. Results

Subjects performed a cued target reaching paradigm while recording MEG using either ‘classic’ SQUID-MEG or a small array of ^4^He-OPM sensors overlying the contralateral sensorimotor area. Data analysis focused on the onset of the reaching target array (‘visual’ epochs) and the subsequent movement, centered on the movement offset (the ‘motor’ epochs).

### 3.1 Time-frequency analysis of motor preparation and execution

The onset of the visual stimulus, and therewith the reaching target was followed by a (typical) reduction in overall beta band power in sensors overlying the sensorimotor area contralateral to the hand, followed by a return back to baseline after the reaching movement. As can be seen in figure 3A, this beta band power reduction was significant in all measured sensor modalities (SQUID-MEG and both OPM-MEG axes), although the extent of the significant frequencies varied, with a larger extent for SQUID-MEG. Recordings using OPM-MEG were more susceptible to low-frequency artefacts during motor execution (right panel), which were maximal at movement offset, causing an artefact that interfered with the beta power rebound measurement. There is an additional significant high frequency (60 – 80 Hz) cluster in the SQUID-MEG data around movement onset, which does not appear in the OPM-MEG data. Figure 3B (right panel) shows that the overall burst rate over time follows the average beta power, although burst rate modulations are clearly weaker. Significant deviations from baseline occur in overall burst rate after onset of the visual cue for SQUID-MEG and for the radial axis of OPM-MEG, although to a smaller extent than the average beta power. The tangential OPM axis does not show a significant modulation of burst rate. For non-baselined burst rate, see supplementary Fig. 1. Unexpectedly, no significant modulations of overall burst rate were found during the ‘motor’ epochs in any modality.

**Figure 3.**
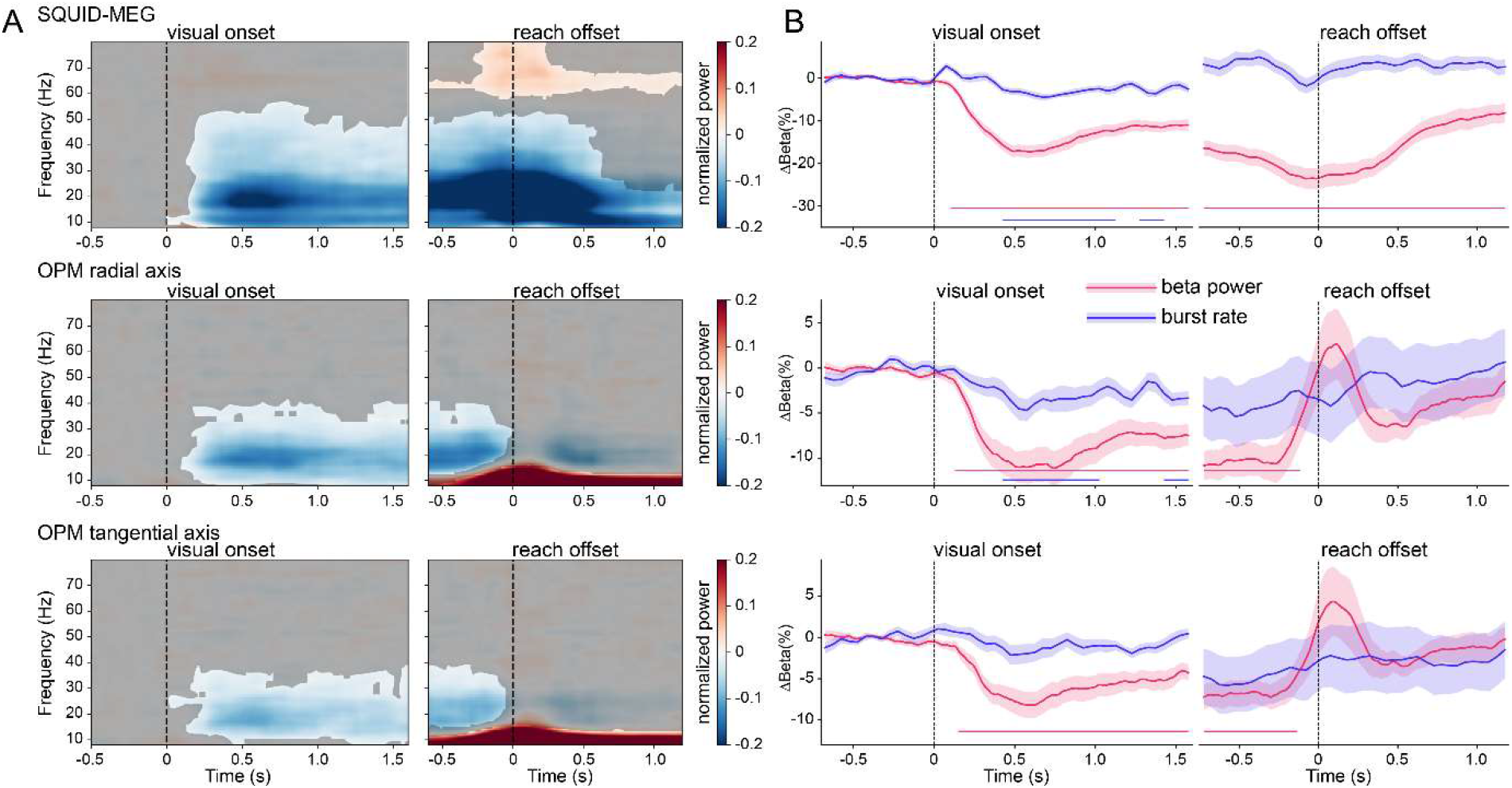
Group averaged power and burst rate. A) Time-frequency representation of the data relative to the onset of visual stimulation and reach offset for SQUID-OPM-radial and OPM-tangential MEG. Non-significant areas are overlayed in gray. B) Burst rate and beta band power over time. Shaded areas denote standard error of the mean. Significant modulations relative to baseline are marked by solid lines at the bottom of the plots.

**Figure 4.**
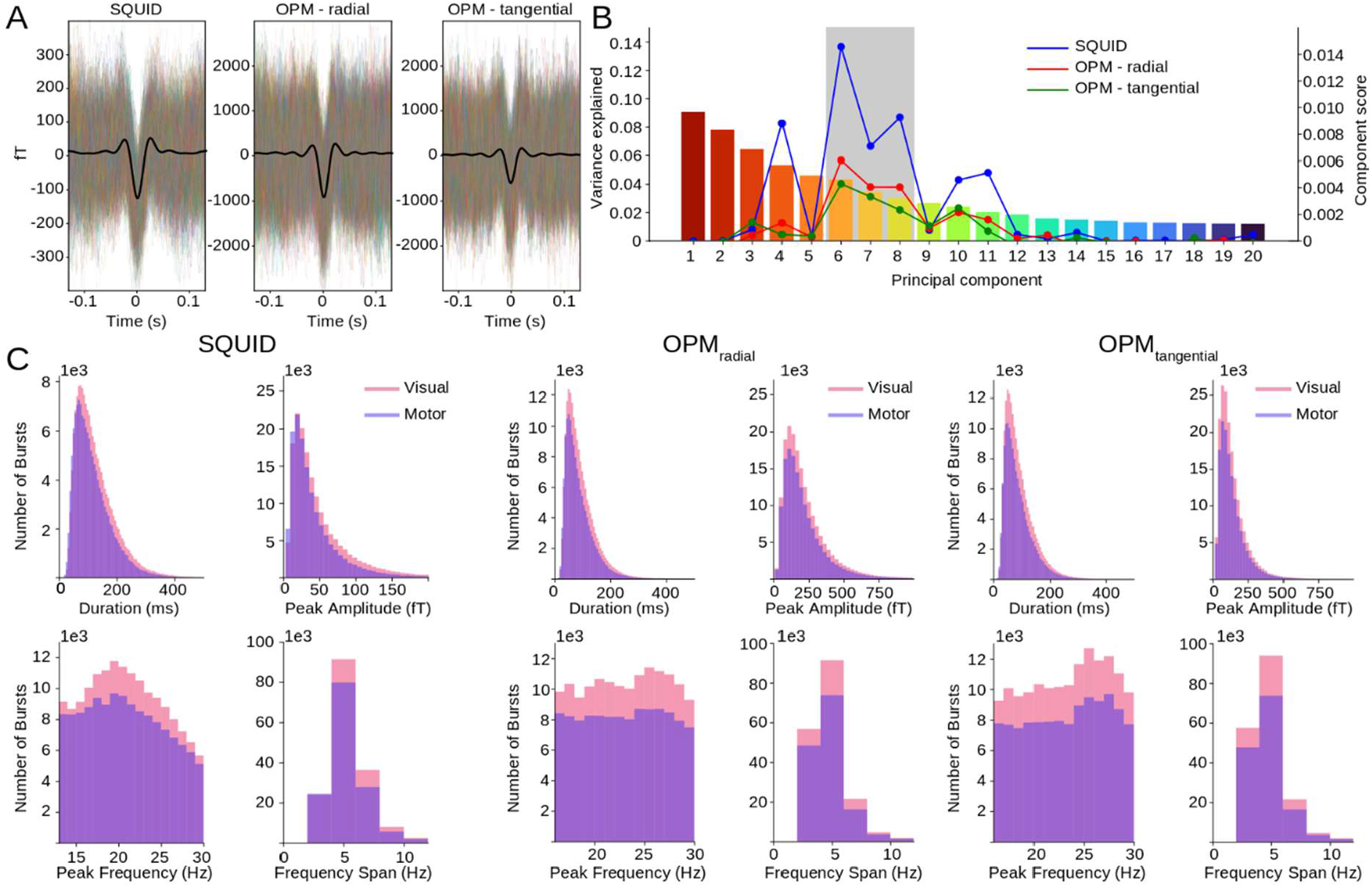
Beta burst metrics. A) Extracted waveforms that formed the basis of the PCA analysis and mean waveform (black) for all modalities. B) PCA components and their explained variance. The colored overlayed lines display the component score used for selection of the PCA components. Selected PCA components are marked in gray. C) Metrics of the extracted waveforms for all modalities (SQUID, OPM) and epochs (visual, motor). Pink = visual, purple = motor).

### 3.2 Beta burst metrics

The beta bursts that were extracted from the data show significant differences between visual and motor trials, as well as between SQUID and OPM data. The average waveform shape (Fig. 4A), however, looks very similar, with a large difference in amplitude (approximately a 5-fold increase), as the OPM-MEG is placed much closer to the scalp, with a resulting increase in signal amplitude, which is most prominent on the radial OPM axis. The metrics extracted from the bursts (Fig. 4C) were also compared, between epochs (‘visual’/’motor’) and measurement modality (‘SQUID’/’OPM’). For brevity, we collapsed across OPM measurement axes, and across epochs for modality comparisons, but full results can be found in the supplemental materials (see supplementary figure 2 and 3). Median values are reported as metrics are generally skewed, as can be seen in Figure 4. Overall, beta bursts were shorter for OPM-MEG than SQUID-MEG (median length 104 ± 66 ms for SQUID-MEG and 76 ± 49 ms for OPM-MEG, D = 0.22, *p* < 0.001). Peak amplitude, as mentioned, was significantly higher for OPM-MEG than SQUID-MEG (median amplitude 33.6 ± 40.1 fT for SQUID-MEG and 141.5 ± 141.5 ft for OPM-MEG, D = 0.66, *p* < 0.001). Peak frequency was lower for SQUID-MEG than OPM-MEG (median frequency 20 ± 4.8 Hz for SQUID-MEG, 22 ± 4.8 Hz for OPM-MEG, D = 0.11, *p* < 0.001). Finally, the frequency span was broader for SQUID-MEG than OPM-MEG, despite identical median values (median frequency span 4 ± 1.83 Hz for SQUID-MEG and 4 ± 1.83 Hz for OPM-MEG, D = 0.17, *p* < 0.001). When comparing between the visual and motor epochs we found longer burst durations for visual epoch bursts than motor epoch bursts (SQUID-MEG - median length visual: 108 ± 69 ms, motor: 100 ± 64 ms, D = 0.052, *p* = 4.7e-181; OPM-MEG - median length visual: 78 ± 50 ms, motor: 74 ± 48 ms, D = 0.043, *p* = 1.2e-250). Peak amplitudes were higher for visual than motor bursts (SQUID-MEG - median amplitude visual: 36.1 ± 43.0 fT, motor: 30.9 ± 35.9 fT, D = 0.081, *p* < 0.001; OPM-MEG - median amplitude visual: 141.9 ± 143.6 fT, motor: 138.7 ± 138,9 fT, D = 0.016, *p* = 1.7e-34. Peak frequency was slightly higher for visual than motor, with equal medians of 20 Hz for SQUID-MEG and 22 Hz for OPM-MEG (SQUID-MEG D = 0.014, p = 3.2e-13; OPM-MEG D = 0.021, p = 2.9e-63. Finally, burst frequency span was slightly broader for visual than motor epochs, despite equal medians of 4 Hz (D = 0.036, p = 4.1e-84; OPM-MEG D = 0.014, p = 1.2e-26. Differences between epochs (visual, motor) were thus significant, though the effects were very small (all D < 0.081).

### 3.3 PCA components of beta bursts

Beta bursts pooled together from all modalities and epochs were subjected to principal component analysis (PCA) to identify dimensions along which burst waveforms vary from the mean waveform shape. The 20 components together explained 63% of the variance in waveform shapes, which may be a little lower due to the heterogeneity of the waveform shapes, as the SQUID-MEG and OPM-MEG waveforms were pooled together. Each burst waveform is thus assigned a score for each PCA component, indicating its place along this dimension (where the center of each dimension is the mean waveform). We selected the components that showed task modulation for further analysis. As the ‘motor’ epochs contained movement related artefacts, PCA components were chosen from the ‘visual’ epochs based on the modulation of burst rate after visual target onset. For comparability, we chose the same three PCA components (components 6,7 & 8) for all modalities (Fig. 4B). A full overview of all components can be found in the supplementary materials (sup. Fig. 4-6) as well as a comparison between waveform shapes extracted from the common PCA and a PCA based on SQUID and OPM data separately (sup Fig. 7). To analyze how specific burst waveforms were modulated over time, the component burst rate was calculated along the entire score range (Fig. 5 left panel) and per binned score quartile (Fig. 5 right panel). Permutation tests were used to determine the significance of the modulations relative to baseline, where non-significant values are masked in the time-dimension 2D plot, and significant values indicated underneath the timecourses on the right panels. This illustrates clearly that bursts with waveforms further away from the mean waveform show larger modulations than those with waveforms closer to the mean. As can be seen in Figure 5 for the SQUID-MEG data, there is a clear and significant reduction in burst rate after visual onset, particularly for the upper extreme percentiles (75-100%), followed by a return to baseline after movement offset. In some cases, the visual onset was even followed by a significant increase in burst rate, as in the first two quartiles (0-25^th^ and 25-50^th^) of PC 6 and the second quartile of PC 7. Interestingly, where the overall burst rate modulation did not reach significance for the *motor* epochs, most bursts with waveforms further away from the mean waveform do show a significant modulation from baseline. These significant modulations are both increases and decreases, which may explain why the overall burst rate is relatively flat, and the effects of interest were occluded. A similar pattern can be observed for the OPM-MEG data. For the radial data, the extent of the significant modulation after visual onset is longer and larger than for the overall burst rate, especially for the upper quartile. The same holds for the tangential axis of the OPM data, although here the overall burst rate does not show any significant modulation, where the specific waveforms do. Unfortunately, the movement artefact also affects the waveform specific burst rates, although significant early modulation (before movement offset) can be seen, which is absent in the overall burst rate. Overall, the specific waveform burst rates, particularly the upper percentile, show the expected pattern of reduction during movement preparation and return to baseline after movement onset. These modulations are clearly stronger than those shown by the overall burst rate.

**Figure 5.**
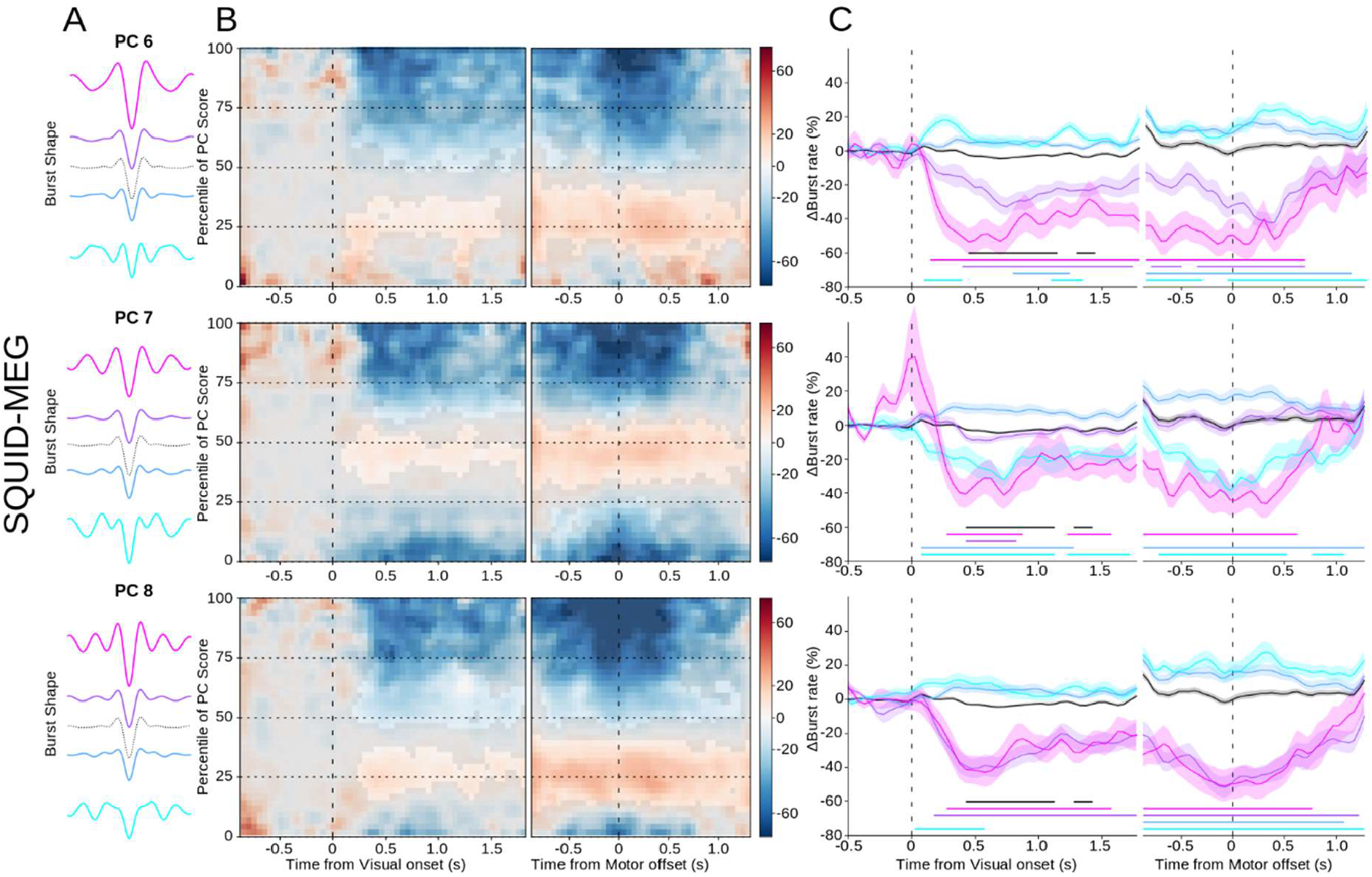
SQUID-MEG specific waveforms. A) Mean waveforms for each quartile of the principal component score range (PCs 6,7 & 8), with the mean overall waveform as a dotted line. B) Mean burst rate over time across the range of scores for the PC component relative to the visual onset (left) or reach offset (right). Non-significant areas are overlayed in gray. C) Burst rate timecourses for bursts with waveforms in each quartile of the PC component, corresponding to the colors in A. Shaded areas denote standard error of the mean. Significant deviations from baseline are marked by solid lines of the corresponding color.

**Figure 6.**
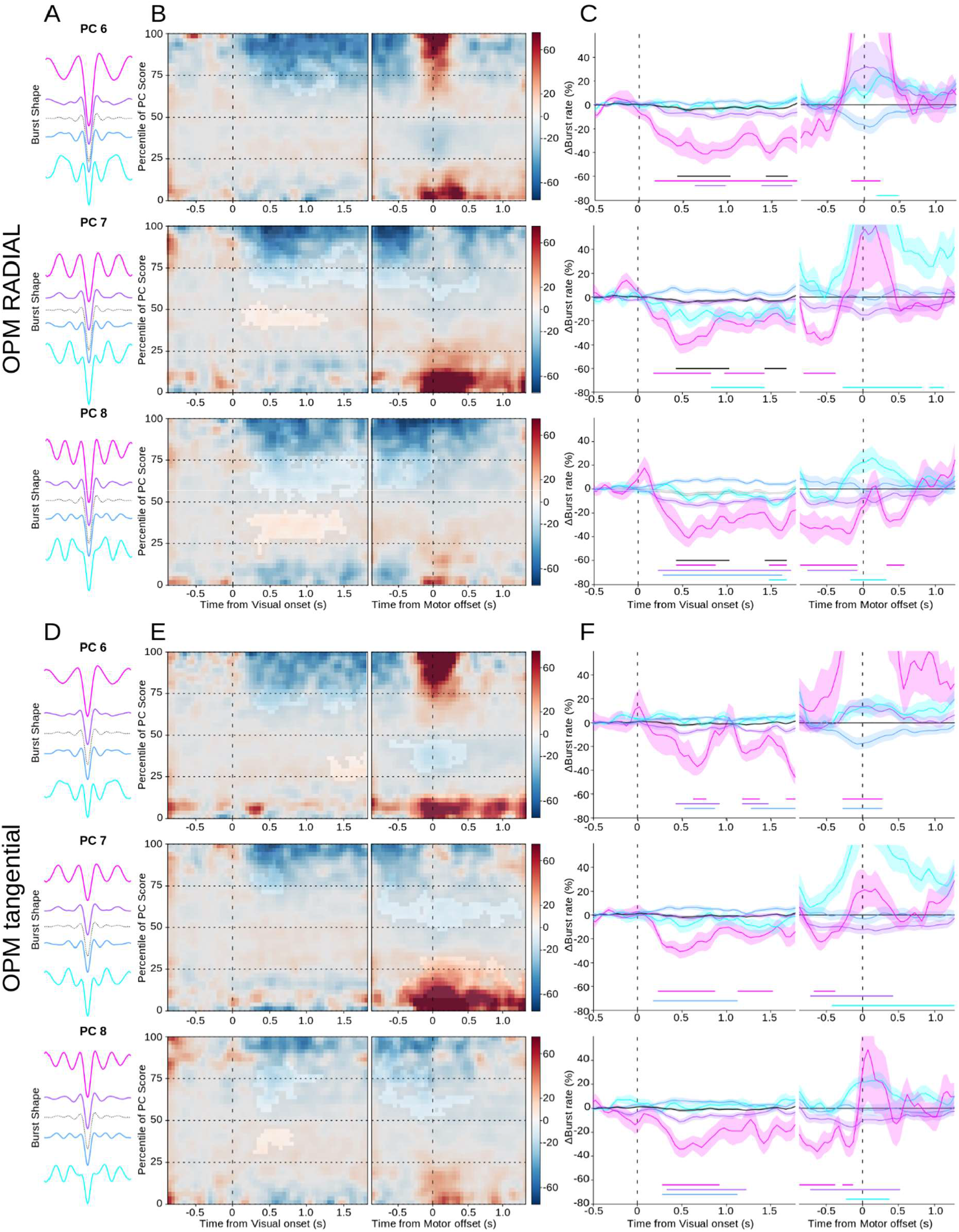
OPM-MEG specific waveforms for the radial (top three) and tangential (bottom three) axes. A) Mean waveforms for each quartile of the score range of the principal component (PCs 6,7 & 8), with the mean overall waveform as a dotted line. B) Mean burst rate over time across the range of PC component scores relative to the visual onset (left) or reach offset (right). Non-significant areas are overlayed in gray. C) Burst rate timecourses for bursts with waveforms in each quartile of the PC component, corresponding to the colors in A. Shaded areas denote standard error of the mean. Significant deviations from baseline are marked by solid lines of the corresponding color. D-E) as A-C for the OPM tangential axis.

### 3.4 SNR

To compare the performance of the ^4^He-OPM sensors to ‘classic’ SQUID-MEG, we computed the signal-to-noise ratio (SNR) of beta-band power. As can be seen in figure 7A, despite the increased amplitude provided by the OPM sensors, SQUID-MEG shows overall higher SNR values than both the radial and tangential OPM axes, but also a much larger range (SQUID mean: 19.4 ± 9.4, OPM_radial_: 8.0 ± 3.8, OPM_tangential_: 5.9 ± 2.7, repeated measures ANOVA: F_(2,34)_: 40.9, p < 0.001). Post hoc tests confirm all sensor types to be different (Wilcoxon signed rank, all comparisons p < 0.001). Note that these calculations were only made on the visual epochs, as the movement artefact in the motor epochs confounds the SNR calculation. Next, we compared data from all sensor types after classification by specific burst type. For selected components (6, 7, 8) the percent baseline change was compared per sensor type and PCA component over subjects (see Fig. 7B). As can be seen, the values here are much closer, with no clear advantage for any sensor type. Results from a repeated measures ANOVA with factor ‘sensor type’ within subjects and ‘PCA component’ between subjects confirm this, as there were no significant results (main effect of sensor type F_(2,102)_ = 2.00, p = 0.141). To quantify the null effect, the same rmANOVA was run using Bayesian statistics. The main effect of sensor shows a Bayes factor BF_01_ = 2.6, indicating the data are 2.6 times more likely under the null hypothesis of no difference.

**Figure 7.**
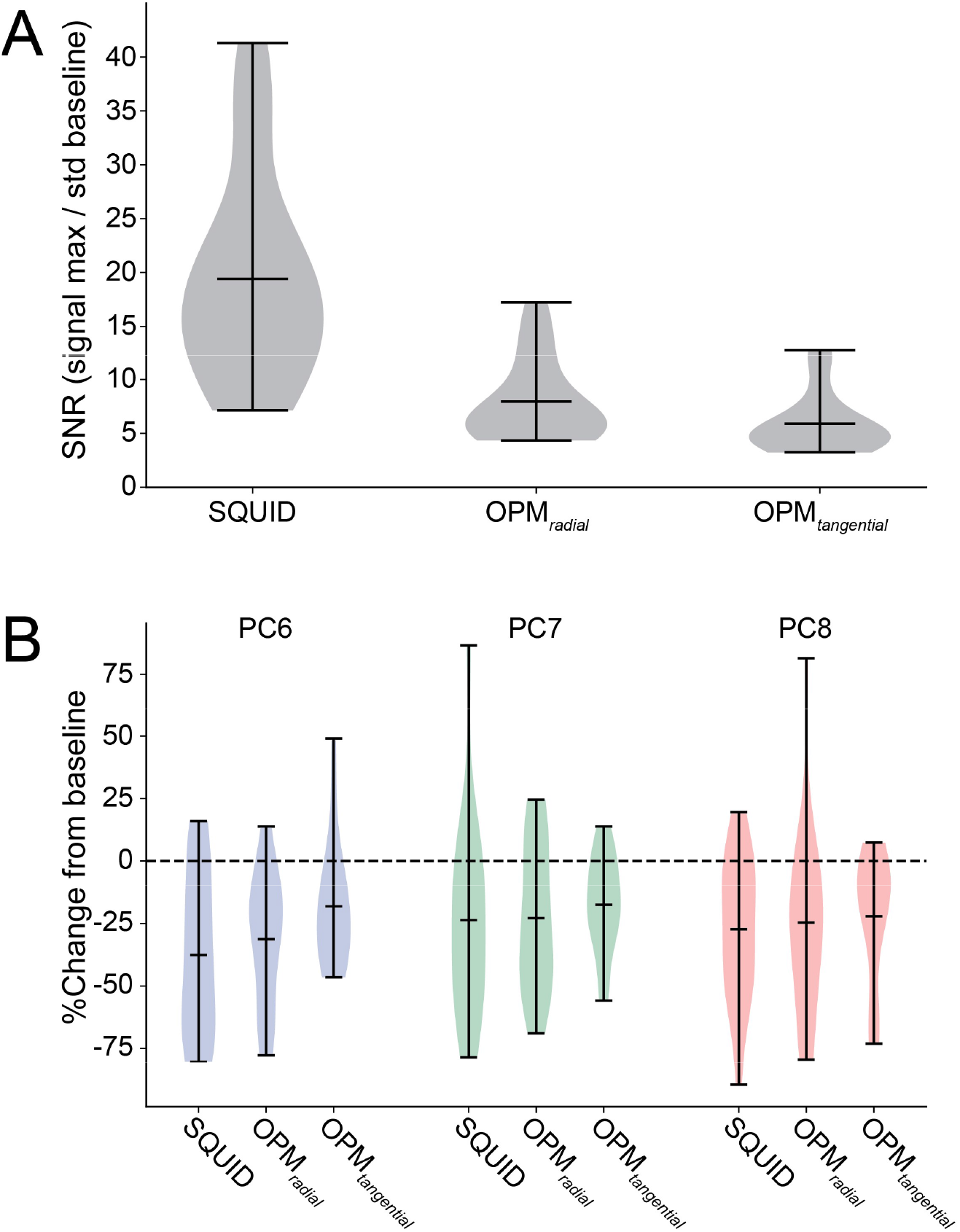
Performance comparison. A) Signal-to-noise ratio calculated on the averaged beta band power for the visual epochs. Central black bars denote the mean. B) Percent change from baseline for the component burst rates, calculated for the quartile with the largest change (the fourth quartile in all cases) for the selected components 6,7 and 8. Central black bars denote the mean.

## 4. Discussion

Here we used a cued visuomotor reaching task to directly compare waveform-specific beta burst dynamics recorded using either ‘classic’ SQUID-MEG or recently developed ^4^He-OPMs. Results show that, while the OPMs are more affected by movement artefacts and a lower SNR in the beta-band range, high quality, task-specific burst waveforms and burst rates could be extracted that were on par with a fully developed SQUID-MEG system.

The time-frequency representation after visual onset of the reach targets revealed a similar spectral pattern for SQUID-MEG and OPM-MEG. While the OPM-MEG data exhibited a somewhat reduced modulation of beta power, there was robust, significant beta range power reduction after the onset of the reach target on both OPM axes. Although occluded somewhat by the movement artefact for the OPM sensors, beta band power returns to baseline after movement cessation, following the expected beta band power pattern (Neuper et al., 2006; Pfurtscheller and Lopes da Silva, 1999), although not exceeding baseline. This may be due to the subsequent movement back to center using the joystick, although this movement was generally effortless. Despite our best efforts, the movement artefact affecting the OPM sensors could not be fully corrected for. In the setup used in this study, it is likely that arm movement had a larger impact on OPM recordings than SQUID recording. This, of course, is the price of free (or less restricted) head movement. Unfortunately, the cable and head support of the OPM system used in this study was not optimal. The cables used were relatively heavy (and these have since been drastically reduced in size), with little support for the head. This may have caused tension in the neck area, which may have compounded (low-frequency) movement related artefacts with more high-frequency components. Additionally, the limited amount of scalp sensors and the single reference sensor do not allow for more sophisticated and promising noise reduction methods that have been developed, such as homogeneous field correction (Tierney et al., 2021), or adaptive multipole modelling (Tierney et al., 2024). Thus, the potential benefit of increased tolerance for movement by using OPMs could not be demonstrated in the current prototype setup. Importantly, while recordings took place in a standard MEG shielded room, there was no advanced (active) shielding. However, at no point during the recording did the OPM sensors saturate or halt data acquisition, but instead remained stable despite large movement artefacts.

When comparing the overall burst waveform, the (wavelet-like) shape is nearly identical for bursts recorded using SQUID-MEG and OPM, although the amplitude is clearly higher for OPM by over a factor four. This is due to both its design, as the SQUID gradiometers have inherently lower amplitude due to their sensitivity to the field *gradient* instead of the absolute field (in contrast to magnetometers), and due to the proximity of the OPM sensors to the scalp and the associated signal increase, in line with expectations (Boto et al., 2018; Iivanainen et al., 2017; Zahran et al., 2022). The latter is likely the main factor, as in a recent study, comparing 4He-OPMs (on scalp) and SQUID-magnetometers (in a standard MEG helmet), the amplitude differed by a factor 2-3 in favor of the scalp OPMs (Badier et al., 2023). Zooming in on the specific metrics, the OPM bursts were also shorter, had a higher peak frequency and a narrower frequency range. As it is unlikely that the underlying brain activity is consistently different, these differences likely arise due to the measurement method. As can be seen from the burst metrics, the peak frequency distribution for SQUID-MEG has a single peak (∼20Hz), whereas the OPM-MEG shows a double peaked distribution (with an additional peak ∼25Hz). This likely also explains the difference in frequency span and length. A previous study using the same method reports a peak frequency around 21Hz (Szul et al., 2023). Thus, it may be that the higher peak frequency may be due to artefactual bursts, possibly due to the OPM system picking up harmonic building vibrations, which peak around 25 Hz on our site. Since the SQUID-MEG system uses gradiometers, these noise sources do not appear in the output signal, but they are picked up by the (OP)magnetometers. Another potential cause of differences between SQUID-MEG and OPM-MEG extracted beta waveform shapes is the differential contribution of superficial and deeper sources. The OPM sensors are magnetometers (sensing far and near) placed directly on the scalp, while the SQUIDs are gradiometers (attenuating far fields) which are placed farther away from the scalp. Given that the magnetic signal drops off exponentially with distance, it is likely that the sensitivity to deeper and superficial sources differ between these sensors. Since the weighing of proximal and distal sources can affect nearly all metrics reported here (Sherman et al., 2016; Szul et al., 2023), peak frequency could also be affected by this. Additionally, and perhaps relatedly, the ^4^He-OPM sensors have three measurement axes, one radial and two tangential. These measure different parts of the magnetic field, and may therefore be sensitive to different neuronal populations than the SQUID sensors.

All extracted bursts, from all modalities, were subjected to principal component analysis and the resulting component dimensions were applied to the sensor specific beta bursts. This yielded very similar waveform shapes for SQUID- and OPM-MEG. Particularly for the upper quartile waveforms, it can be noted that the waveform contains clear additional deflections around the burst center. These seem to drop off a little slower for the OPM derived waveforms. The modulations of waveform specific burst rate also follow the same pattern for both modalities. Interestingly, similarly to the observation in Szul (2023), some waveforms increase their burst rate after visual onset (e.g. the second quartile of PC 8). These waveforms seem to be shorter (with less deflections). This challenges the notion that there is a single homogeneous beta ‘burst’ shape. Instead, the waveform shape is variable over time, where some burst types even increase during action preparation. Interestingly, some waveform shapes were more susceptible to the movement artefact in the OPM data. Perhaps unsurprisingly, lower frequency waveforms (e.g. PC 6) seemed more susceptible than those with a higher frequency (e.g. PC 8). Thus, it may simply be that the low frequency movement artefact correlated more with low frequency waveforms, causing an apparent ‘increase’ in burst rate.

Modulations of overall beta burst rate were more modest than average beta band power, although the burst rate timecourse generally followed the same pattern as the power, as would be expected. However, the waveform specific burst rates, particularly the top quartiles, show much larger modulations over time. As mentioned, this may be due to the teasing apart of waveforms increasing and decreasing the burst rate during over time, which would be occluded in the overall burst rate.

We found that the signal-to-noise ratio of the OPM sensors, as measured from the beta band power was lower than for SQUID-MEG. This is consistent with previous results using these sensors, although the SNR values for ERPs have been found to be closer (Gutteling et al., 2023). However, comparing the burst rate modulations for the selected principal components, no differences were found between SQUID and OPM-MEG. This not only illustrates the strength of the burst detection and waveform analysis used, but also that data quality obtained from (^4^He)OPM sensors can be equal or better than conventional SQUID-MEG. It is important to note that in the meantime, a new generation of ^4^He-OPM sensors have been developed that have a sensitivity better than 30 ft/√Hz, as compared to the ∼45 ft/√Hz used in the current study.

The results in the current study are encouraging, especially given the limitations of the ^4^He-OPM setup used. The setup consisted of only four sensors on the scalp, with a single reference sensor on top of the helmet, and no active shielding. We used a relatively simple processing pipeline, based on regression and ICA. This was compared with a fully developed SQUID-MEG system array of 275 gradiometers and 29 reference sensors. Additionally, as mentioned above, the version of the ^4^He-OPM system used to record the current data had relatively large, inflexible cables which likely compounded head movement of the subject with sensor movement due to the stiff cables. At time of writing, these issues have been resolved and a whole-head system with thin, flexible cables, an improved mounting system and improved sensitivity has been developed. As a future direction, we hope to assess and exploit the benefits that this system provides, and take advantage of higher sensor count and lower noise floor to have increased movement tolerance, lower noise and higher spatial precision. Overall, this is an encouraging step towards unrestricted MEG recordings, opening up new avenues of research using more naturalistic movement paradigms as well as novel avenues for BCI applications (Papadopoulos et al., 2024).

## Data and Code Availability

Anonymized data supporting the results of this study are available from the corresponding author upon reasonable request and validation by regulatory and ethical bodies and subject consent.

Main analysis code can be found here: https://github.com/tgutteling/OPM_Motor

Beta burst extraction pipeline: https://github.com/danclab/burst_detection

### Author Contributions

*Conceptualization*: J.J.B., D.S., E.L., J.J. *Methodology*: T.P.G., E.L., D.S., J.J.B, M.J.S. *Software*: T.P.G., M.J.S., J.J.B., D.S., E.L. *Formal analysis*: T.P.G. *Investigation*: D.S., J.J.B, M.J.S, E.L., S.D. *Data curation*: D.S. W*riting—original draft preparation*: T.P.G., E.L., J.J.B., D.S. *Writing—review and editing*: T.P.G., M.J.S., S.D., E.L., J.J., J.J.B., D.S. *Project administration*: D.S., E.L., J.J. *Funding acquisition*: D.S., E.L., J.J.

All authors have read and agreed to the published version of the manuscript

### Funding

This project was funded by Région Auvergne-Rhône-Alpes (Pack Ambition project: NEW_MEG). This project was partly funded by France Life Imaging (WP2 & 4) (grant “Infras-tructure d’avenir en Biologie Santé” ANR-11-INBS-0006). TG is supported by the Labex Cortex (ANR-11-LABX-0042).

M.J.S and J.J.B. were supported by the European Research Council under the European Union’s Horizon 2020 research and innovation program (ERC consolidator Grant 864550 to J.J.B).

### Declaration of Competing Interests

The authors declare the following competing interests: E.L. holds founding equity in Mag4Health SAS, a French start-up company that is developing and commercializing MEG systems based on He-OPM technology. Mag4Health SAS provided technical support to the data acquisition. For the recordings performed until 1 February 2022, E.L. was still an employee of CEA LETI.

## Notes

https://github.com/tgutteling/OPM_Motor

