## Supplementary materials for "Beta burst waveform extraction using novel ^4^He-OPMs"

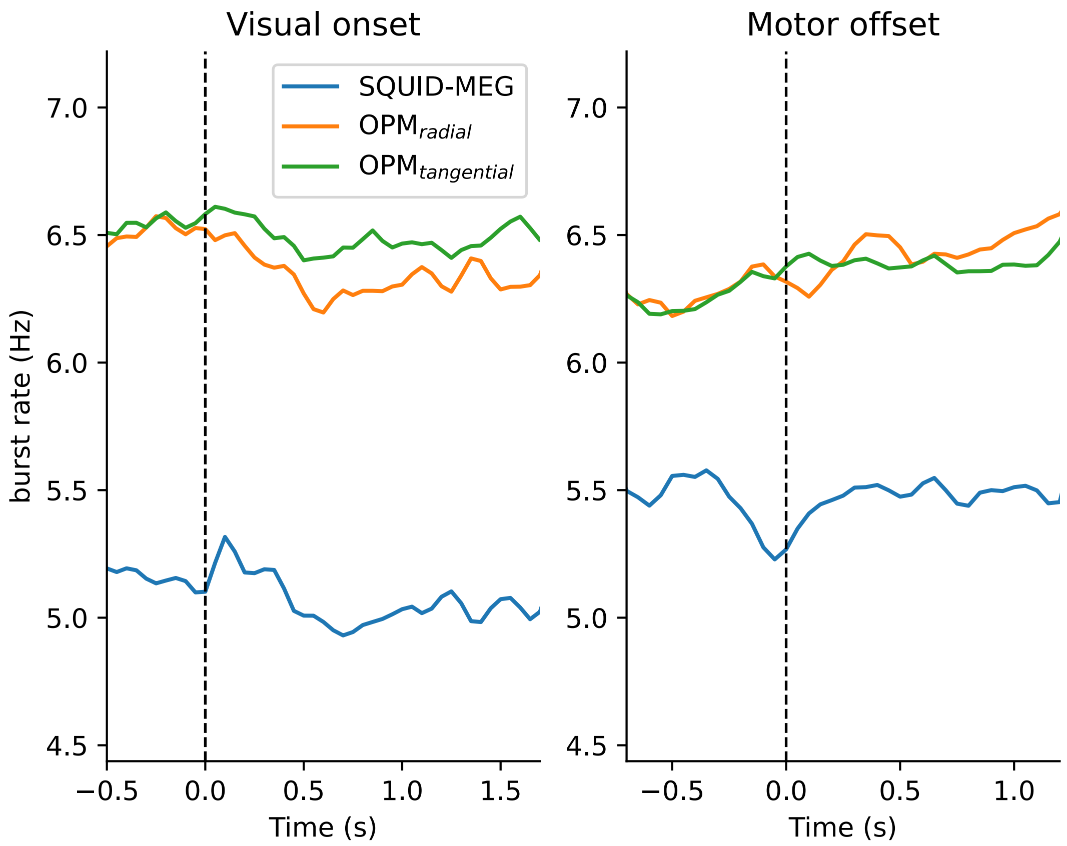


Supplementary figure 1 – Non-baselined beta burst rates for SQUID-MEG and OPM radial and tangential axes. Burst rate is expressed as the average individual bursts per second, per trial, per sensor.

| Metric | Modality | Mean | Median | SD |
|---|---|---|---|---|
| Duration (ms) | SQUID | 124.58 | 108.00 | 68.61 |
| | OPM rad | 90.74 | 78.00 | 50.68 |
| | OPM tan | 90.05 | 78.00 | 49.85 |
| Peak Amplitude (fT) | SQUID | 50.00 | 36.11 | 43.05 |
| | OPM rad | 224.14 | 179.27 | 166.31 |
| | OPM tan | 140.34 | 114.15 | 101.04 |
| Peak Frequency | SQUID | 20.74 | 20.00 | 4.76 |
| | OPM rad | 21.93 | 22.00 | 4.78 |
| | OPM tan | 22.16 | 22.00 | 4.77 |
| Frequency Span | SQUID | 4.53 | 4.00 | 1.85 |
| | OPM rad | 3.85 | 4.00 | 1.85 |
| | OPM tan | 3.83 | 4.00 | 1.81 |

Supplementary figure 2 – Means, medians and standard deviations for all metrics extracted from the beta bursts.

### A Condition tests
Visual vs. motor epochs

| Metric | Modality | D | P |
|---|---|---|---|
| Duration (ms) | SQUID | 0.053 | 5.94E-182 |
| | OPM rad | 0.049 | 1.30E-169 |
| | OPM tan | 0.036 | 8.63E-93 |
| Peak Amplitude (fT) | SQUID | 0.081 | 0.00E+00 |
| | OPM rad | 0.025 | 8.13E-44 |
| | OPM tan | 0.011 | 9.02E-09 |
| Peak Frequency | SQUID | 0.014 | 4.03E-14 |
| | OPM rad | 0.023 | 3.35E-37 |
| | OPM tan | 0.020 | 5.24E-28 |
| Frequency Span | SQUID | 0.036 | 5.12E-85 |
| | OPM rad | 0.015 | 5.27E-16 |
| | OPM tan | 0.013 | 4.44E-12 |

### B Modality tests
Visual epochs

| **Duration** | SQUID | OPMrad | OPMtan |
|---|---|---|---|
| SQUID | | 0.224 | 0.229 |
| OPMrad | 0.224 | | 0.005 |
| OPMtan | 0.229 | 0.005 | |

| **Peak amplitude** | SQUID | OPMrad | OPMtan |
|---|---|---|---|
| SQUID | | 0.718 | 0.572 |
| OPMrad | 0.718 | | 0.268 |
| OPMtan | 0.572 | 0.268 | |

| **Peak frequency** | SQUID | OPMrad | OPMtan |
|---|---|---|---|
| SQUID | | 0.109 | 0.129 |
| OPMrad | 0.109 | | 0.025 |
| OPMtan | 0.129 | 0.025 | |

| **Frequency span** | SQUID | OPMrad | OPMtan |
|---|---|---|---|
| SQUID | | 0.173 | 0.172 |
| OPMrad | 0.173 | | 0.004 |
| OPMtan | 0.172 | 0.004 | |

### C Modality tests
Motor epochs

| **Duration** | SQUID | OPMrad | OPMtan |
|---|---|---|---|
| SQUID | | 0.219 | 0.209 |
| OPMrad | 0.219 | | 0.011 |
| OPMtan | 0.209 | 0.011 | |

| **Peak amplitude** | SQUID | OPMrad | OPMtan |
|---|---|---|---|
| SQUID | | 0.767 | 0.639 |
| OPMrad | 0.767 | | 0.257 |
| OPMtan | 0.639 | 0.257 | |

| **Peak frequency** | SQUID | OPMrad | OPMtan |
|---|---|---|---|
| SQUID | | 0.118 | 0.118 |
| OPMrad | 0.118 | | 0.032 |
| OPMtan | 0.118 | 0.032 | |

| **Frequency span** | SQUID | OPMrad | OPMtan |
|---|---|---|---|
| SQUID | | 0.161 | 0.158 |
| OPMrad | 0.161 | | 0.003 |
| OPMtan | 0.158 | 0.003 | |

Supplementary figure 3 – Full Kolmogorov-Smirnov tests results. A) Tests between conditions (visual epochs vs motor epochs) for the burst metrics extracted and imaging modalities. B) Tests comparing metrics between modalities for visual epochs and C) motor epochs. All $p<.022$ except when indicated in red.

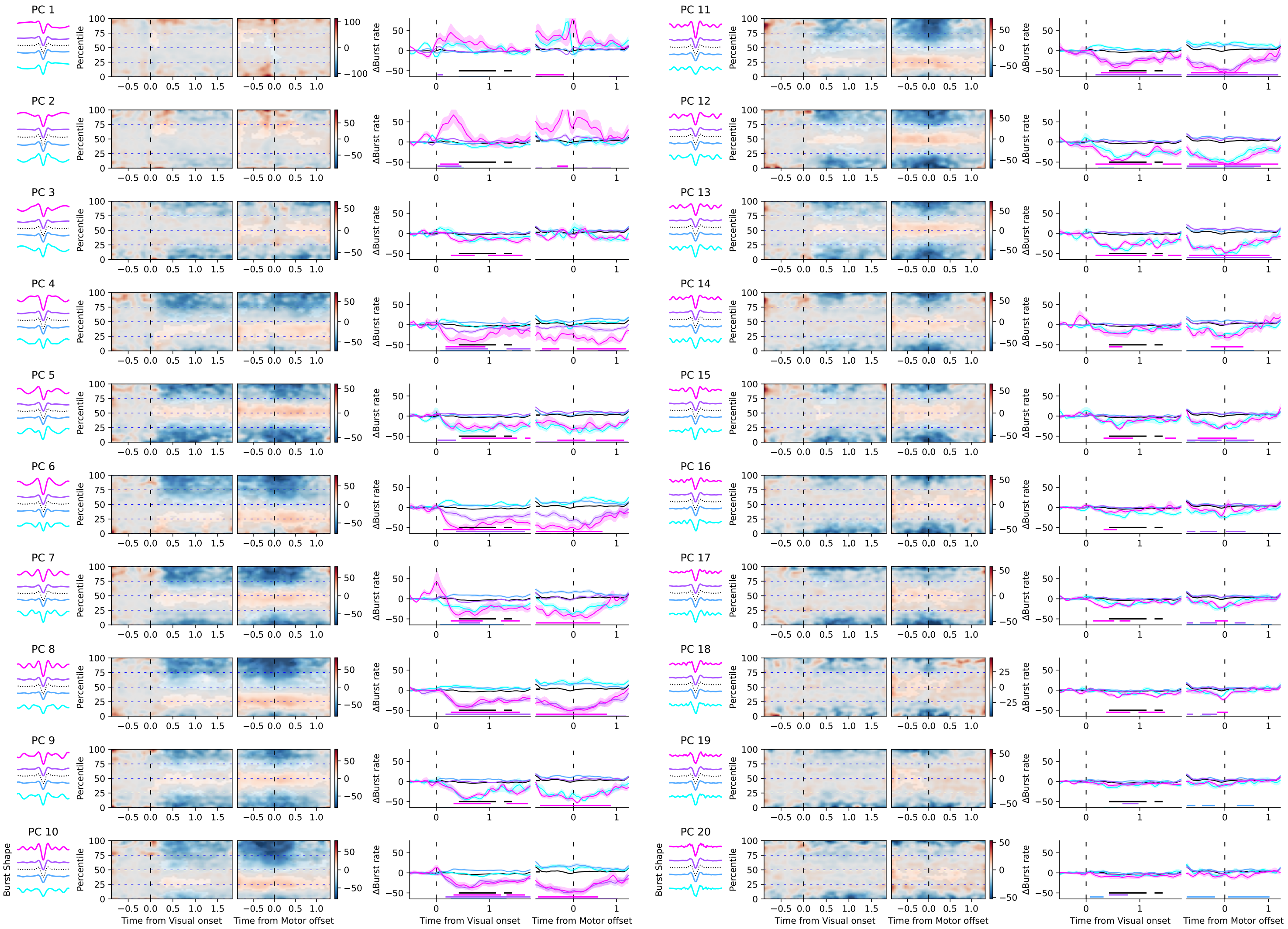


Supplementary figure 4 – SQUID-MEG specific waveforms for all principal components. Per component from left to right: Mean waveforms for each quartile of the principal component score range, with the mean overall waveform as a dotted line; Mean burst rate over time across the range of scores for the PC component relative to the visual onset (left) or reach offset (right), with non-significant areas are overlayed in gray; Burst rate timecourses for bursts with waveforms in each quartile of the PC component. Significant deviations from baseline are marked by solid lines of the corresponding color.

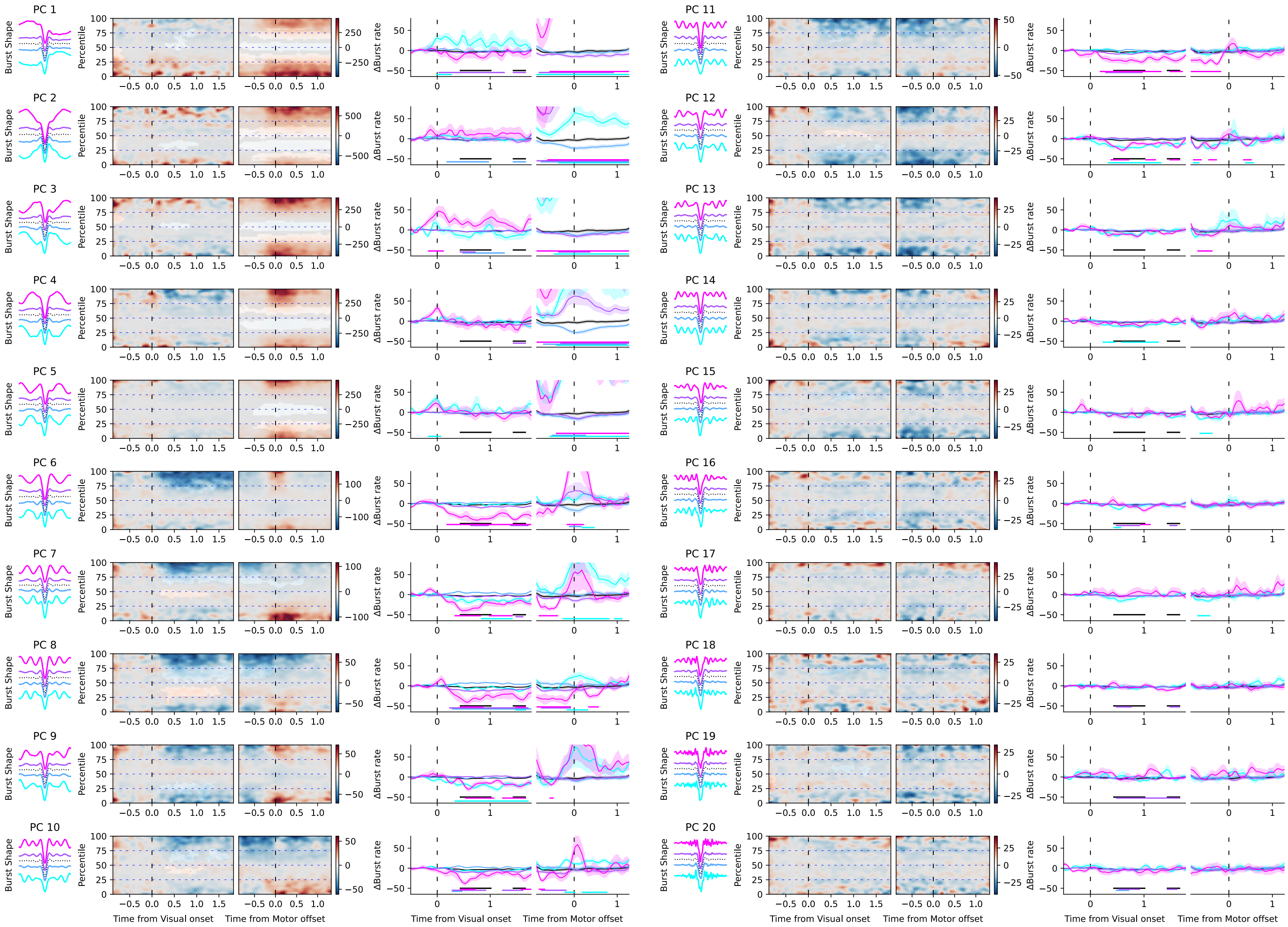

Supplementary figure 5 – OPM - radial axis specific waveforms for all principal components. As figure 3, per component from left to right: Mean waveforms for each quartile of the principal component score range, with the mean overall waveform as a dotted line; Mean burst rate over time across the range of scores for the PC component relative to the visual onset (left) or reach offset (right), with non-significant areas are overlayed in gray; Burst rate timecourses for bursts with waveforms in each quartile of the PC component. Significant deviations from baseline are marked by solid lines of the corresponding color.

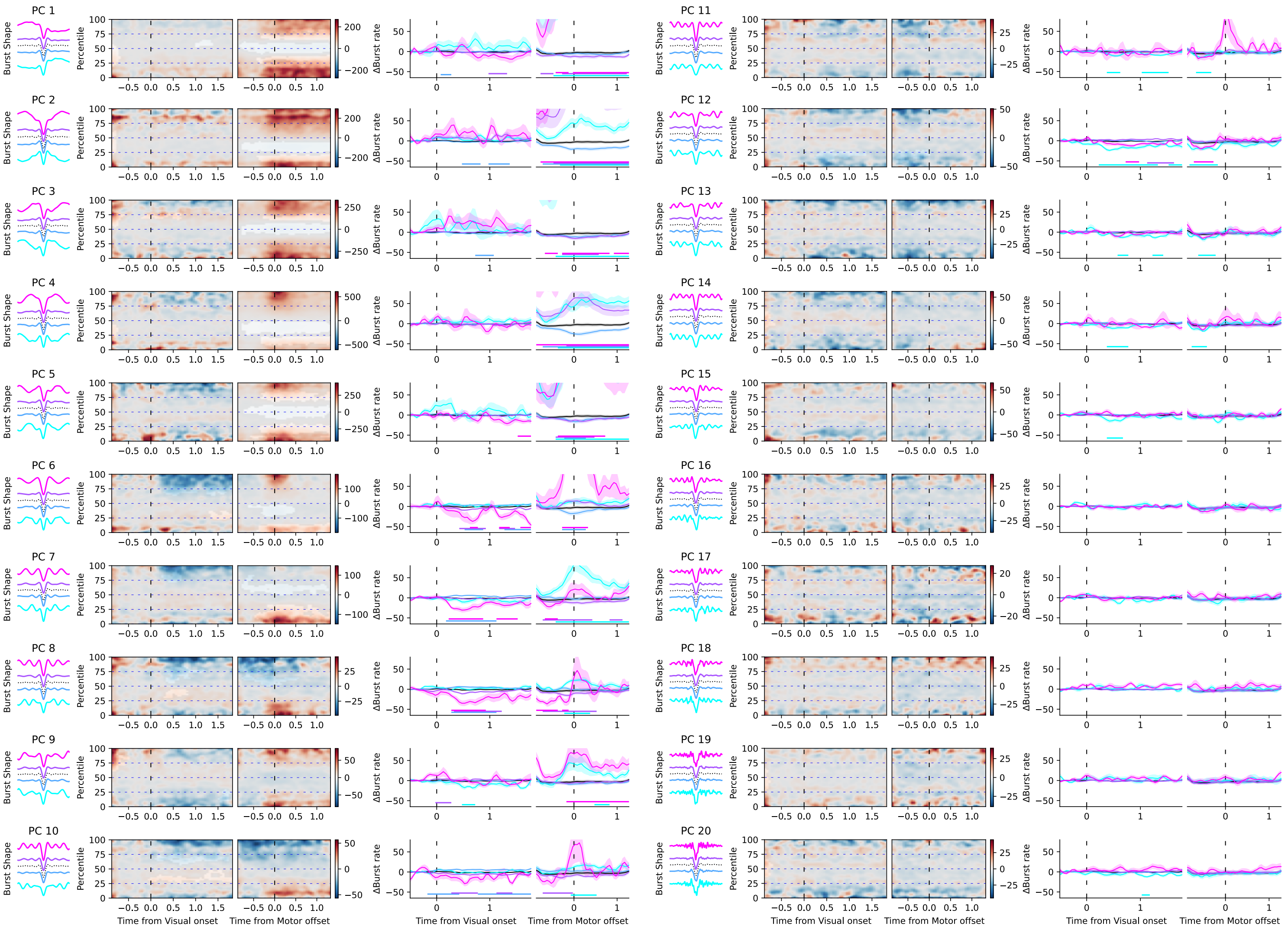


Supplementary figure 6 – OPM - tangential axis specific waveforms for all principal components. As figure 3-4, per component from left to right: Mean waveforms for each quartile of the principal component score range, with the mean overall waveform as a dotted line; Mean burst rate over time across the range of scores for the PC component relative to the visual onset (left) or reach offset (right), with non-significant areas are overlayed in gray; Burst rate timecourses for bursts with waveforms in each quartile of the PC component. Significant deviations from baseline are marked by solid lines of the corresponding color.

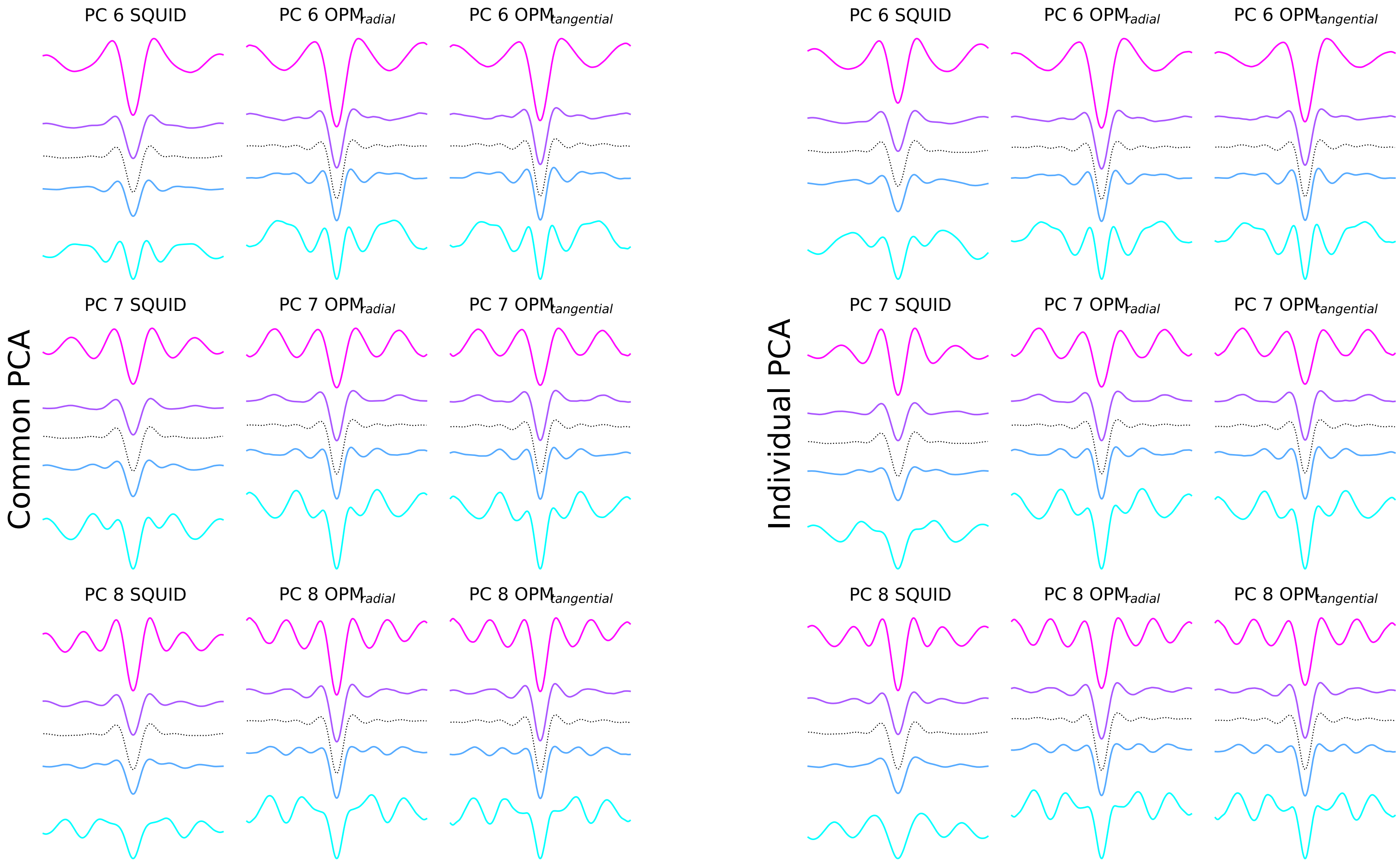


Supplementary figure 7 – Comparison of specific waveforms resulting from either a common PCA (left), pooling all burst together from SQUID-MEG and OPM data and an modality specific 'individual' PCA (right), based on bursts derived either from SQUID-MEG data (left column) or the radial and tangential OPM axes.

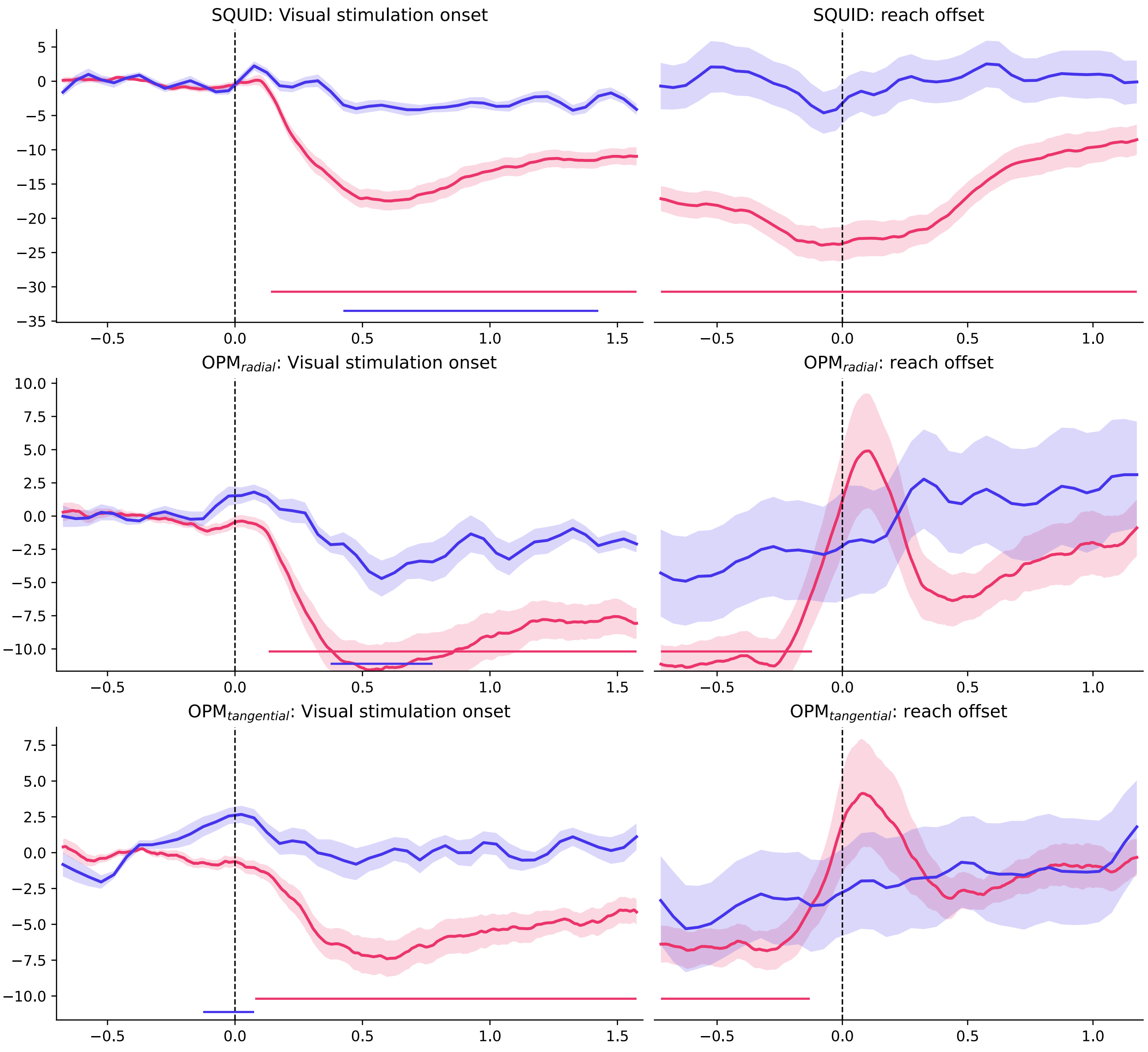


Supplementary figure 8 - Group averaged power and burst rate derived from data preprocessed without ICA. As main Fig. 3B. Burst rate and beta band power over time. Shaded areas denote standard error of the mean. Significant modulations relative to baseline are marked by solid lines at the bottom of the plots. Red line: beta power; Blue line: burst rate.

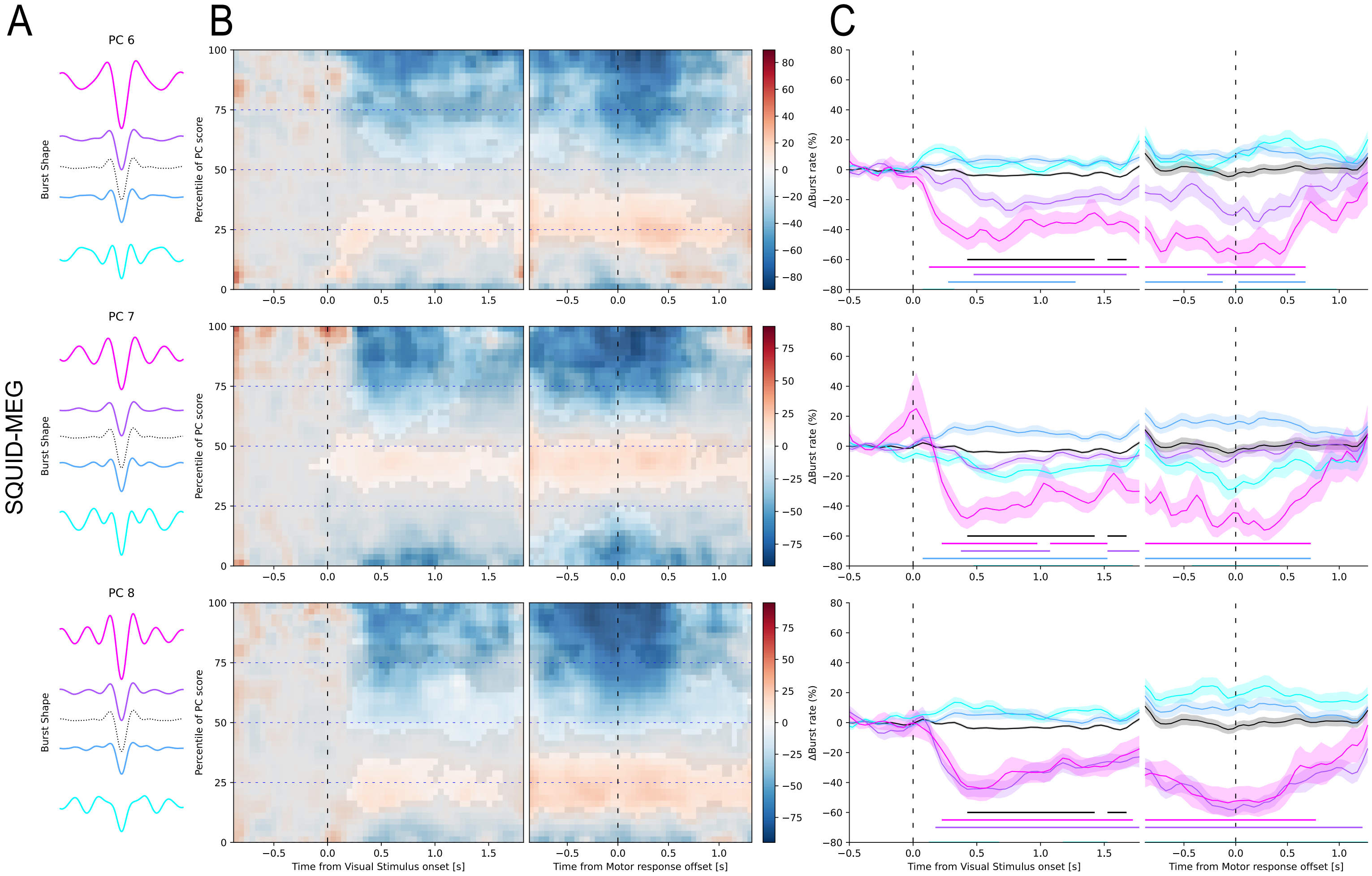


Supplementary figure 9 - SQUID-MEG specific waveforms derived from data preprocessed without ICA. As main Fig. 5. A) Mean waveforms for each quartile of the principal component score range (PCs 6,7 & 8), with the mean overall waveform as a dotted line. B) Mean burst rate over time across the range of scores for the PC component relative to the visual onset (left) or reach offset (right). Non-significant areas are overlayed in gray. C) Burst rate timecourses for bursts with waveforms in each quartile of the PC component, corresponding to the colors in A. Shaded areas denote standard error of the mean. Significant deviations from baseline are marked by solid lines of the corresponding color.

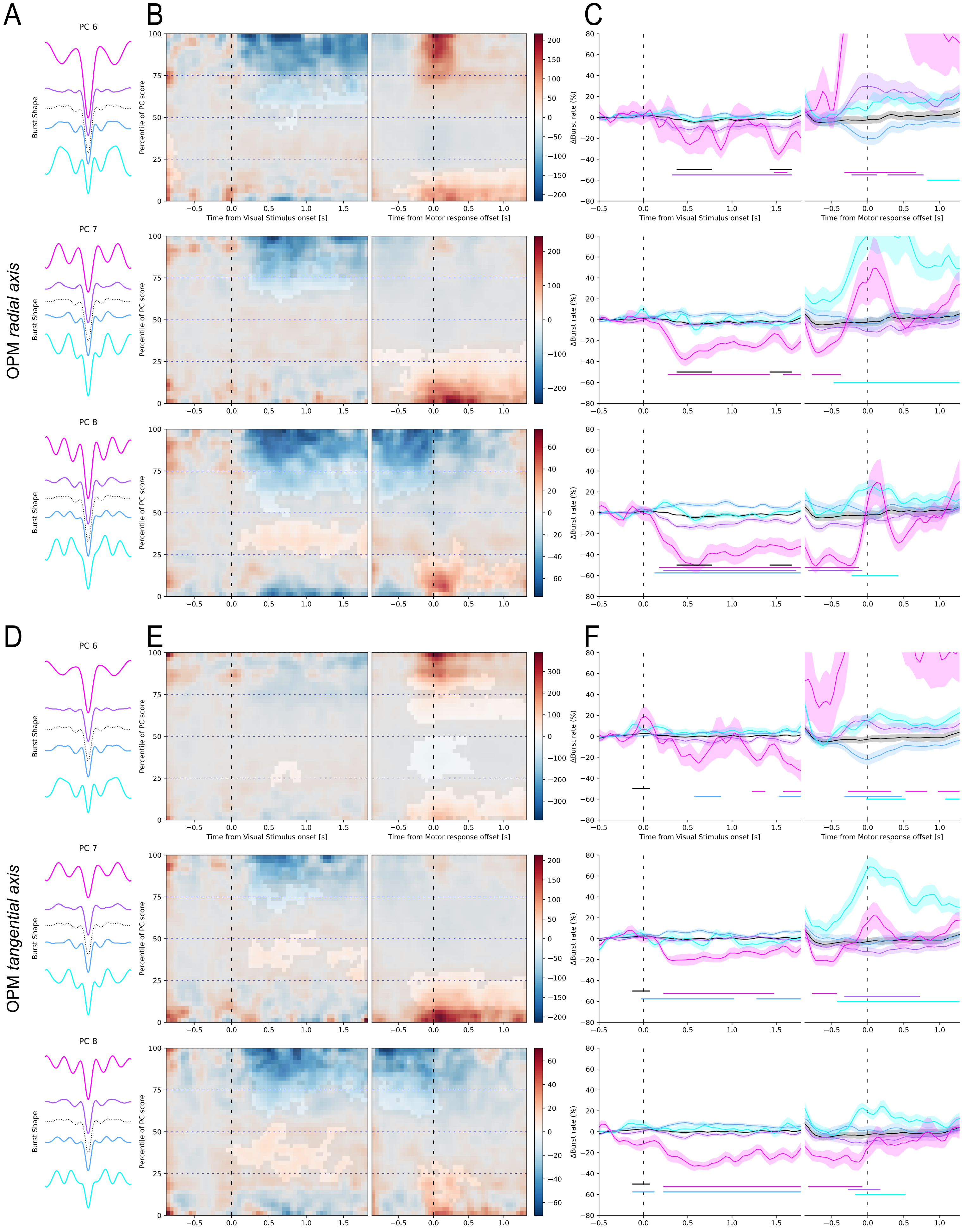


Supplementary figure 10 - OPM specific waveforms derived from data preprocessed without ICA. As main Fig. 6. A) Mean waveforms for each quartile of the principal component score range (PCs 6,7 & 8), with the mean overall waveform as a dotted line. B) Mean burst rate over time across the range of scores for the PC component relative to the visual onset (left) or reach offset (right). Non-significant areas are overlayed in gray. C) Burst rate time-courses for bursts with waveforms in each quartile of the PC component, corresponding to the colors in A. Shaded areas denote standard error of the mean. Significant deviations from baseline are marked by solid lines of the corresponding color. D-E) as A-C for the OPM tangential axis.
